# Usiigaci: Instance-aware cell tracking in stain-free phase contrast microscopy enabled by machine learning

**DOI:** 10.1101/524041

**Authors:** Hsieh-Fu Tsai, Joanna Gajda, Tyler F.W. Sloan, Andrei Rares, Amy Q. Shen

## Abstract

Stain-free, single-cell segmentation and tracking is tantamount to the holy grail of microscopic cell migration analysis. Phase contrast microscopy (PCM) images with cells at high density are notoriously difficult to segment accurately; thus, manual segmentation remains the de facto standard practice. In this work, we introduce Usiigaci, an all-in-one, semi-automated pipeline to segment, track, and visualize cell movement and morphological changes in PCM. Stain-free, instance-aware segmentation is accomplished using a mask regional convolutional neural network (Mask R-CNN). A Trackpy-based cell tracker with a graphical user interface is developed for cell tracking and data verification. The performance of Usiigaci is validated with electrotaxis of NIH/3T3 fibroblasts. Usiigaci provides highly accurate cell movement and morphological information for quantitative cell migration analysis.

## 1. Motivation and significance

Cell migration is a fundamental cell behavior that underlies various physiological processes, including development, tissue maintenance, immunity, and tissue regeneration, as well as pathological processes such as metastasis. Many *in vitro* as well as *in vivo* platforms have been developed to investigate molecular mechanisms underlying cell migration in different microenvironments with the aid of microscopy. To analyze single-or collective-cell migration, reliable segmentation of each individual cell in microscopic images is necessary in order to extract location as well as morphological information.

Among bright-field microscopy techniques, Zernike’s phase contrast microscopy (PCM) is favored by biologists for its ability to translate phase differences from cellular components into amplitude differences, so as to make the cell membrane, the nucleus, and vacuoles more visible [1]. However, PCM images are notoriously difficult to segment correctly using conventional computer vision methods, due to the low contrast between cells and their background [2]. For this reason, many cell migration experiments still rely on fluorescent labeling of cells or manual tracking. Fluorescent labeling of cells requires transgenic expression of fluorescent proteins or cells tagged with fluorescent compounds, both of which can be toxic to cells and which require extensive validation of phenotypic changes. Although thresholding fluorescent images is relatively straightforward, cells that are in close proximity are often indistinguishable in threshold results. On the other hand, manual tracking of cell migration is labor-intensive and prone to operator error. Conducting high-throughput microscopy experiments is already possible thanks to methodology and instrumental advances, but current analytical techniques to interpret results quantitatively face major obstacles due to imperfect cell segmentation and tracking [3]. Moreover, cell movement is not the only parameter of interest in cell migration. For cell migration guided by environmental gradients, shear stress, surface topology, and electric field can also impact cell morphology [4–7].

Although many software packages have been developed for cell tracking, the majority of them handle only fluorescent images and require good thresholding results [8]. While some software tackles stain-free cell tracking, outlining each individual cell accurately to the cell boundary is difficult; thus, these packages are limited to positional tracking and cannot resolve adjacent or touching cells [8–12]. Migrating cells in ameboid or mesenchymal mode often have thin protruding cellular structures for locomotion, such as blebs or lamellipodia [13]. These structures exhibit very low contrast in PCM, which prevents reliable segmentation, even though they are essential for cell migration.

In recent years, advances in machine learning using convolutional neural networks (CNNs) have proven effective at solving computer vision problems [14–16]. Among them, Deepcell architecture, proposed by Van Valen *et al*., has demonstrated that cells in close proximity can be segmented using pixel-wise classification of the background, the cell membrane, and the cell cytoplasm [15]. However, fluorescent staining of cell nuclei is still needed for optimal segmentation of these PCM images.

To address the above challenges, we introduce newly developed stain-free, instance-aware cell tracking software for PCM, called Usiigaci. Stain-free, instance-aware segmentation of phase contrast microscopy images is appealing to biologists because cells are free of labeling damage and their analysis does not suffer from false readings. Moreover, both locations and outlines of cells can be analyzed in their entirety.

## 2. Software description

### 2.1. Software overview

Usiigaci, pronounced as *ushi:gachi* by Hepburn romanization, is a Ryukyuan word that refers to tracing the outlines of objects, which is an appropriate description of the function of our software. Usiigaci has a semi-automated workflow consisting of three modules: a segmentation module, a tracking module, and a data processing module, all written in standard Python syntax (Figure 1).

**Figure 1:**
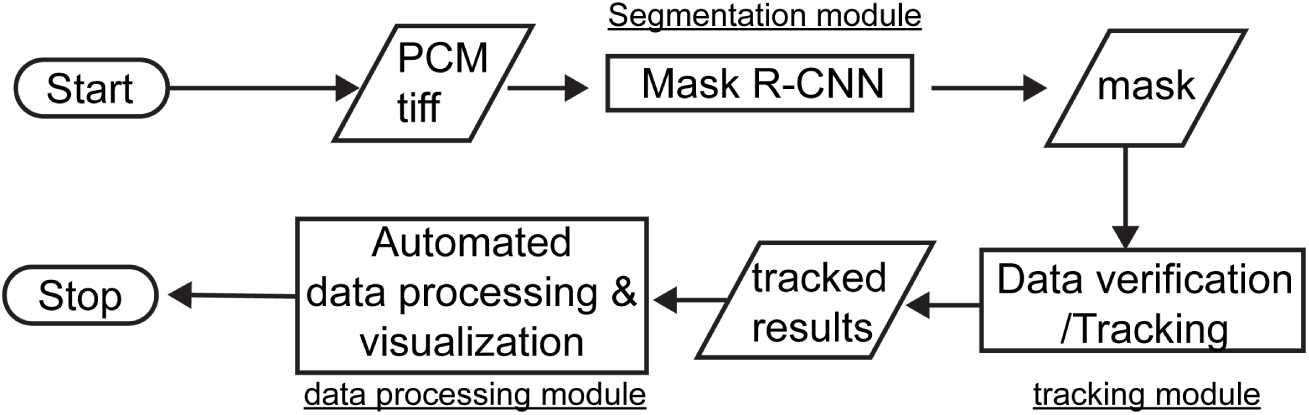
The all-in-one segmentation, tracking, and data processing workflow of Usiigaci.

A Mask R-CNN model pretrained with the Microsoft COCO dataset [17] was further trained using 50 manually annotated PCM images with single cell outlines as a classification class (more details in S1.4 and S1.5 in the SI document for preparing custom training data and to initiate new training). Using this trained model, PCM images are provided as input to the Mask R-CNN-based segmentation module and highly accurate instance-aware segmented masks are generated [18]. Outlines of individual cells in the images are correctly segmented into identifiers (IDs), even if they are in close proximity. IDs are then linked and tracked in the tracking module. With the aid of a graphical user interface (GUI) in the tracking module, side-by-side comparison of PCM images and tracked masks allow users to validate segmentation and tracking results. At this point, unwanted cell tracks, such as imperfectly segmented or tracked cells, mitotic cells, or dead cells can be excluded by users prior to data processing. Thereafter, step-centric and cell-centric parameters of cell migration, as well as visualization of cell migration data are computed and generated automatically from the tracked results in the data processing module (Table S.1 in the SI document).

Based on the three modules described above, Usiigaci is an all-in-one, semi-automated solution for stain-free cell migration analysis in PCM, with a biologist-friendly workflow.

### 2.2. Software architecture and functionality

A diagram of segmentation and tracking modules of Usiigaci is shown in Figure 2. The segmentation module of Usiigaci is based on a Mask R-CNN model that is implemented in TensorFlow and Keras, as originally open-sourced by Matterport Inc. under the MIT license [19–21]. A detailed diagram of Mask R-CNN architecture is shown in Fig. S.3 in the SI document. The Mask R-CNN model is built upon the Faster R-CNN model that has achieved rapid identification of objects through searching regions of interest (ROIs) on feature maps [18, 22]. Raw images undergo multiple convolutional operations in a R-CNN backbone, which is composed of a residual function network (ResNet-101, [23]) and a feature pyramid network (FPN, [24]), to generate 5 feature maps (C1 to C5). ROIs are searched on feature maps using region proposal layers. An accurate instance-segmented ROI map is generated by an ROI align layer to correct for misalignment in the ROIPooling operation. After upsampling, entire outlines of individual cells are segmented into polygons bearing unique IDs in the exported mask. As a result, highly accurate, instance-aware segmentation of stain-free PCM images is realized.

**Figure 2:**
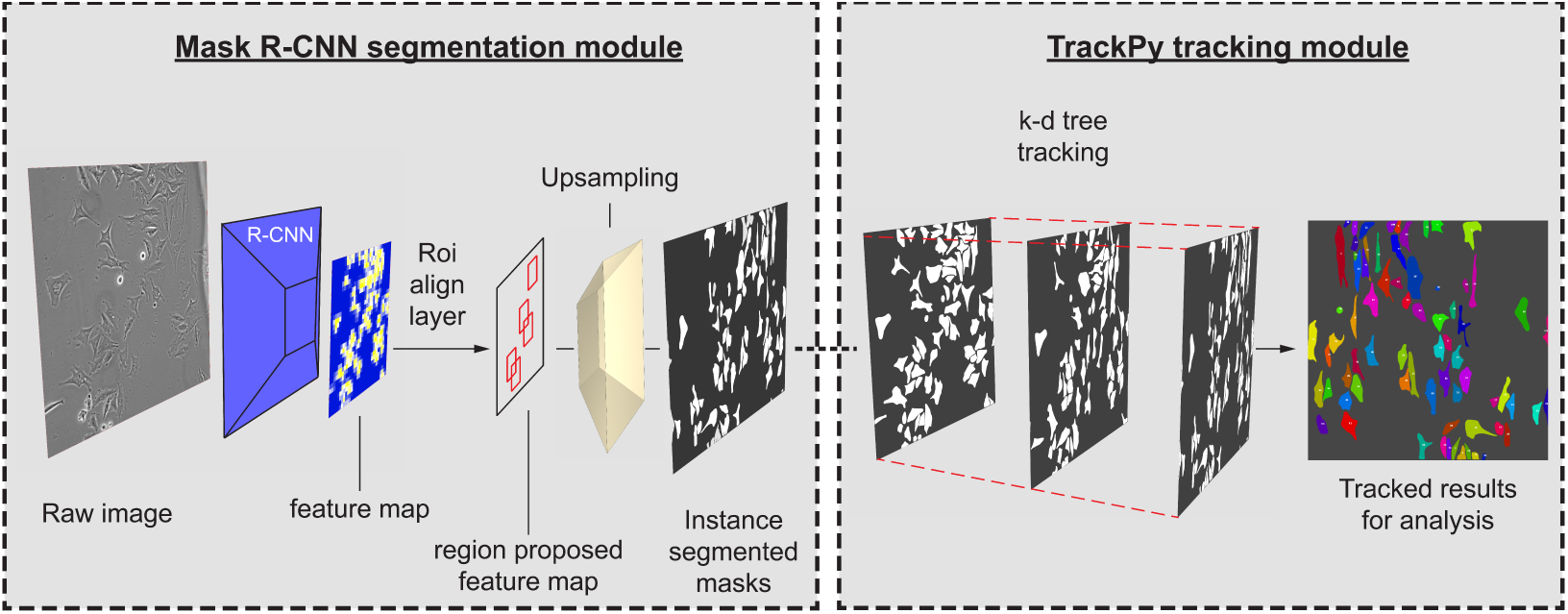
Diagram of segmentation and tracking modules of Usiigaci. PCM images are processed in a Mask R-CNN segmentation module with a region proposal network, which has a backbone of ResNet-101 and a feature pyramid network (FPN), to generate instance-segmented masks. Objects in the masks are linked and tracked in a Trackpy-based tracker using the k-dimensional tree algorithm. Important cell migration parameters are then computed from the tracked results.

After segmentation, each mask contains segmented cell outlines bearing a unique identifier (ID). The IDs are then used for linking and tracking in the tracking module built on the Trackpy library [25]. The features of an ID, such as location, equivalent diameter, perimeter, eccentricity, orientation, and true solidity, are used as parameters in Trackpy for tracking. IDs in each consecutive mask in a time-lapse experiment belonging to the same cell are searched by the Trackpy library using its default nearest neighbor search option, namely the k-dimension tree algorithm [26–28].

Linking and tracking are followed by automatic post-processing, where segmentation and tracking results are corrected in two steps. In the first step, a cell wrongly segmented as two IDs is corrected by merging the two IDs. In the second step, IDs in consecutive frames belong to the same track, but suffering from interrupted events are re-linked. A GUI based on the PyQt and PyQtGraph library for the tracking module is developed so that users can verify segmentation and tracking results [29, 30]. Manual verification is important because imperfections in segmentation can cause errors in tracking. In addition, cells that undergo mitosis and cells that enter or exit the viewfield during the experiment generate tracking results that are not meaningful in single cell migration studies (Fig. S.6 in the SI document). In the GUI of the tracking module, by imposing a simple criterion, *select complete tracks*, the valid tracks IDs of which exist in every frame, can be selected. Thereafter, users can manually verify whether the tracking is correct by cross-referencing against raw images. The amount of labor in the proposed workflow is less than that associated with conventional manual tracking [4]. Subsequently, centroid and morphology parameters such as angle, perimeter, and area of each ID in valid tracks can be extracted and produced using the scikit-image library [31].

Analysis of single-cell migration data is accomplished in the data processing module to compute migration parameters for each ID throughout the time-lapse experiment (Fig. S.1.B). Several data processing libraries, including the Python data analysis library (Pandas), NumPy, and SciPy, are used for processing cell migration data [32–34]. Step-centric and cell-centric features, such as turning angle, net trigonometric distance, speed, orientation, and directedness are computed automatically in a Jupyter Notebook (Table S.1) [35, 36]. Moreover, automated visualization of cell migration in cell trajectory plots, box plots, and time-series plots is generated with the aid of Matplotlib and Seaborn plotting libraries (Fig. S.9) [37, 38].

## 3. Validation of Usiigaci

### 3.1. Segmentation module

Stain-free tracking of NIH/3T3 fibroblasts electrotaxis in a 300 V/m direct current electric field (dcEF) for 10 hr under PCM is used to demonstrate unique features of Usiigaci. Details of cell experiments and imaging are described in the supplementary information. Segmentation and tracking performance of Usiigaci is benchmarked against state-of-the-art free software such as PHANTAST [11], Fogbank [12], Deepcell [15] as well as proprietary software such as Imaris and Metamorph. Segmentation results of Usiigaci and aforementioned software are shown in Figure 3 and quantitatively analyzed by segmentation evaluation metrics (Table S.2 in the SI document). Segmentation similarity can be evaluated using the mean ratio of intersection over union (mIoU), which is also known as the Jaccard index (Figure 4).

**Figure 3:**
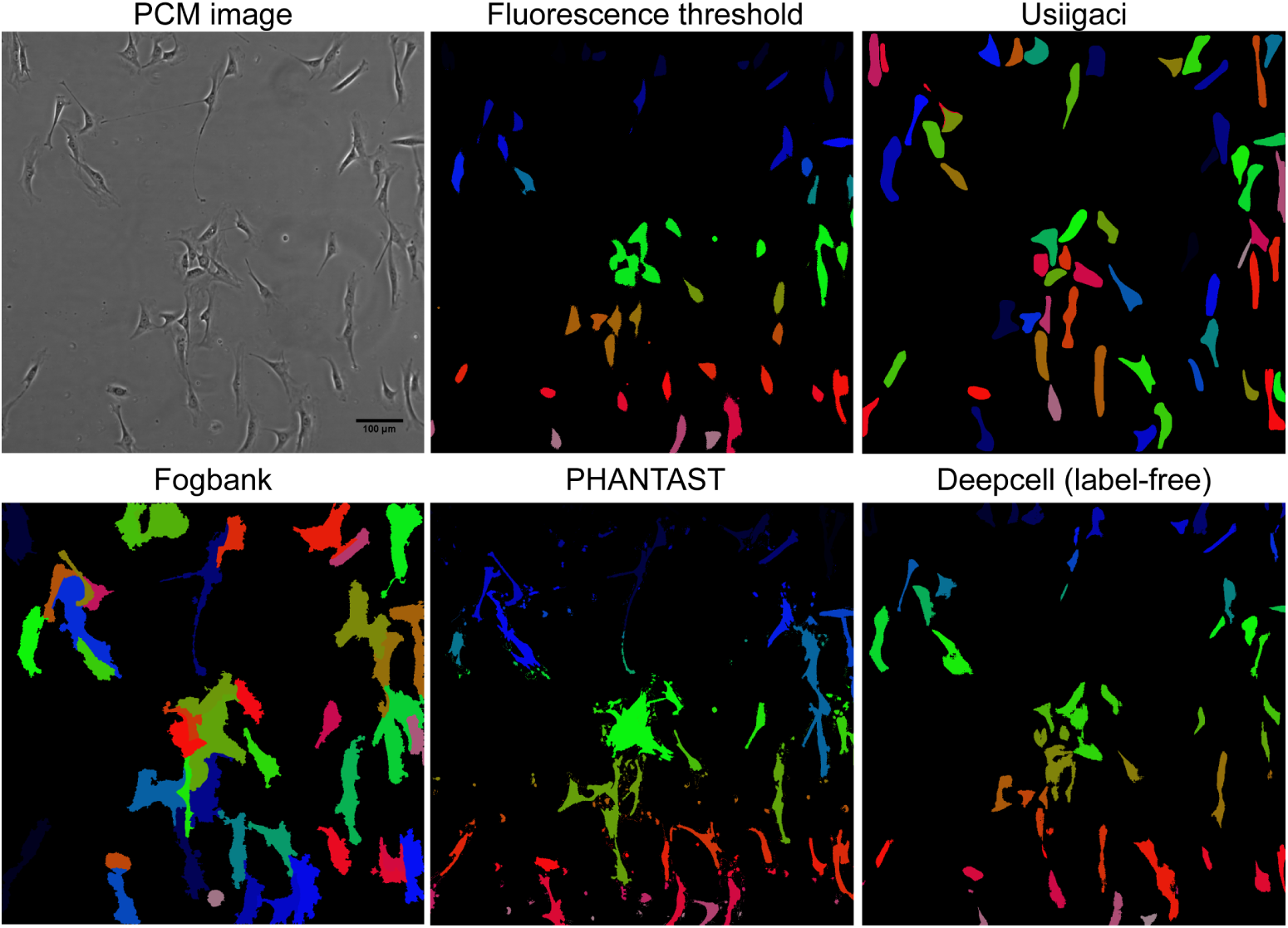
Microscopy of NIH/3T3 cells stained with CellTracker Green under PCM and fluorescence microscopy, compared with segmentation results of Usiigaci, Fogbank, PHANTAST, and Deepcell on the PCM image. Different color represents instances of each region of interest. In Usiigaci, Fogbank, and Deepcell, each cell is segmented into an instance outline with a unique ID and color. In segmented masks of fluorescence-thresholded or PHANTAST, cells are segmented into ROIs using the *analyze particle* function in ImageJ and filled with pseudocolors using the ROImap function in the LOCI plug-in. Usiigaci accurately segmented each individual cell with accuracy superior to that of other software.

**Figure 4:**
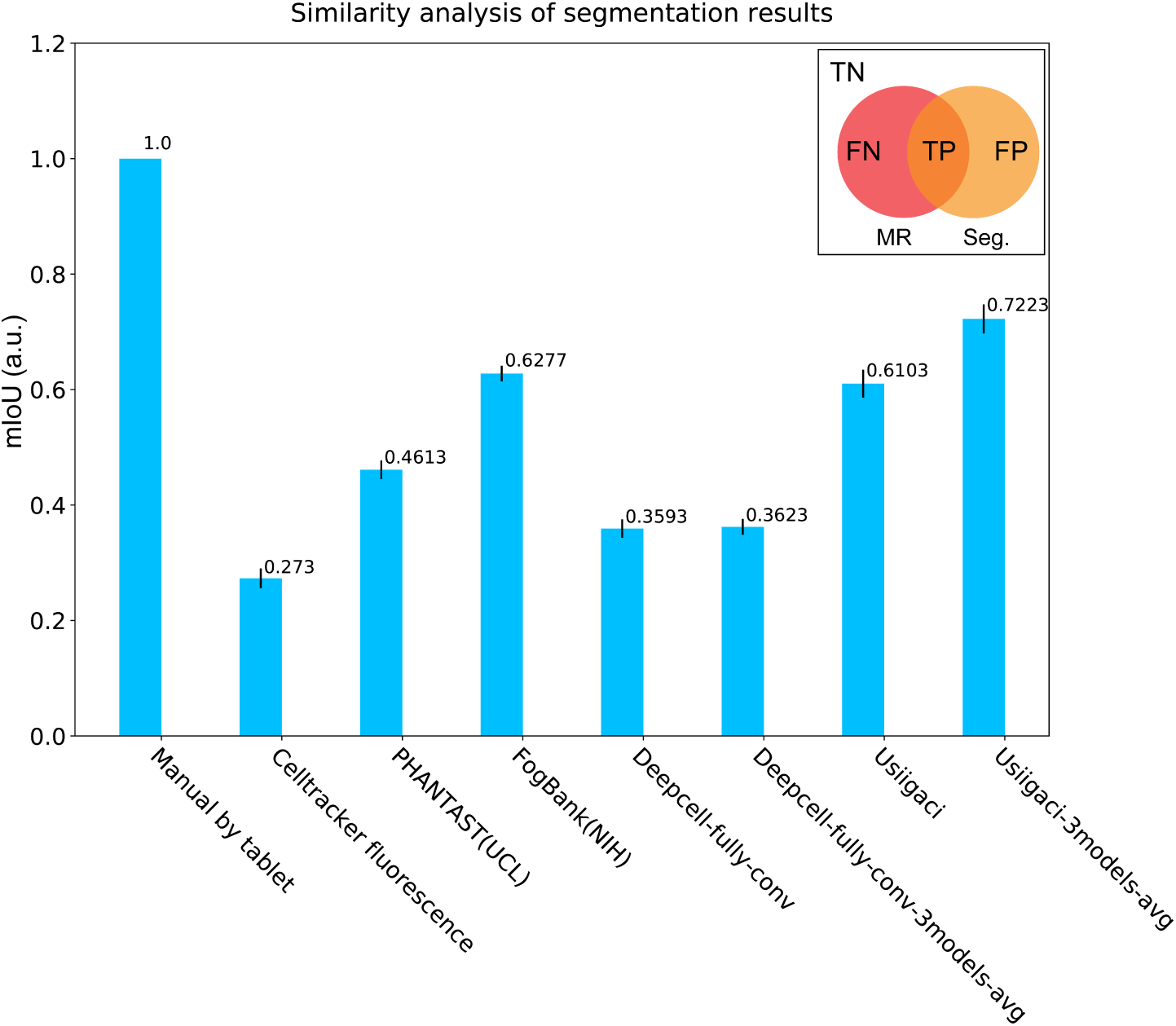
Segmentation similarity averaged among three NIH/3T3 cell images using various methods. MR: Manual reference; Seg.: Segmented results; FN: False negative; TP: True positive; FP: False positive; TN:True negative. Segmentation similarity is measured by the mean intersection over union between ground truth and segmented results 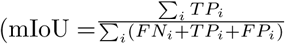, as shown in the inset), or also known as the Jaccard index.

By fluorescence thresholding, thicker cell bodies can be segmented easily, but thinner structures, such as lamellipodia or blebs, often fail to be segmented and contribute to higher specificity and lower mIoU (Table 1 & Fig. S.7 in the SI document). In Fogbank and PHANTAST, images are thresholded by local contrast, thus segmentation is effective only if single cells are well isolated. The segmentation similarity achieved by Fogbank and PHANTAST is moderately high (mIoU 0.46 and 0.63), but single-cell tracking in images with high cell density is not effective using these two methods, because individual cells cannot be distinguished. By classifying cell membranes through machine learning methods, Deepcell segments high density cells better than conventional methods. However, due to the pixel-level classification methods in Deepcell, adjacent cells without clear boundaries are sometimes difficult to segment. In Usiigaci, entire outlines of cells are segmented correctly in an instance-aware fashion, even if cells are densely packed. The segmentation similarity of Usiigaci with a single trained model is 2.2 times higher than that of the fluorescence threshold method. Usiigaci’s segmentation also outperforms other benchmarked segmentation software (Table 1 & Figure 4). Moreover, the segmentation speed of Usiigaci is fast in comparison to manual segmentation and benchmarked software (see Fig. S.8 in the SI document).

**Table 1:** Segmentation performance averaged among three NIH/3T3 cell images using various methods. Ch:channel

|  | Ch | Jaccard index | F1 score | Precision | Recall | Specificity | Accuracy |
| --- | --- | --- | --- | --- | --- | --- | --- |
| Manual | PCM | 1 | 1 | 1 | 1 | 1 | 1 |
| Automatic threshold | FL | 0.27±0.03 | 0.46±0.02 | 0.97±0.02 | 0.30±0.01 | 1±0 | 0.91±0.01 |
| PHANTAST | PCM | 0.46±0.02 | 0.59±0.09 | 0.70±0.19 | 0.51±0.02 | 0.97±0.03 | 0.91±0.03 |
| Fogbank | PCM | 0.63±0.02 | 0.77±0.02 | 0.65±0.02 | 0.93±0.02 | 0.94±0.01 | 0.94±0.01 |
| Deepcell 3models-avg | PCM | 0.36±0.04 | 0.56±0.06 | 0.39±0.06 | 0.96±0.01 | 0.92±0.01 | 0.92±0.01 |
| Usiigaci 3models-avg | PCM | 0.72±0.01 | 0.85±0.01 | 0.83±0.02 | 0.87±0.01 | 0.95±0.04 | 0.96±0.01 |

However, the potential limitation of Usiigaci’s Mask R-CNN (essentially a machine learning method), is that segmentation accuracy may be profoundly impacted if the segmentation image is significantly different from that in the training dataset (see detailed discussion in section S2.3 in the SI document). A proper training dataset created by end users with a user-specific experimental configuration may be necessary for optimal results. The detailed description of training data preparation and training process in supplementary section S1.4 and S1.5 should help users to achieve optimal results if a new training dataset is required.

### 3.2. Tracking module

Mask R-CNN segments cells in an instance-aware manner such that each segmented cell possesses a unique ID (shown with pseudo-color in Figure 3).

The IDs in consecutive images are linked and tracked in the tracking module. A GUI is developed to provide manual data verification for users to identify potential errors in segmentation and tracking (Figure 5). A simple criterion, *select complete tracks*, is built in the GUI for selecting tracks with IDs that exist in every frame. Imposing the criterion ensures high probability of valid tracks (Fig. S.6). Furthermore, the validity of cell tracks can be verified by users. Tracks that are biologically invalid, such as those having cells that have undergone mitosis or cell death, can be excluded manually. Usiigaci provides a labor-saving workflow while preserving the capacity for human intervention, which is essential to ensure data validity in single-cell migration analysis [39].

**Figure 5:**
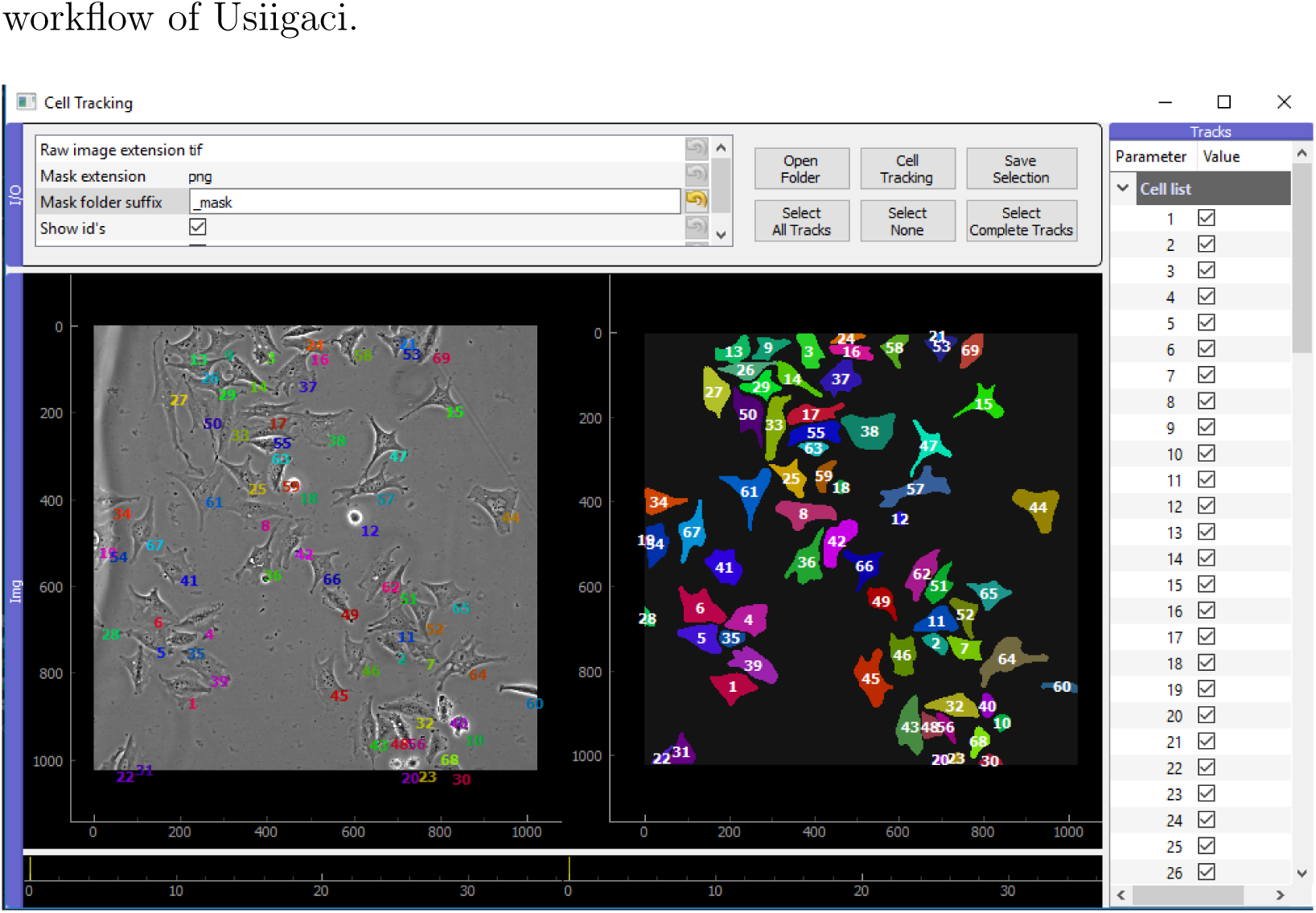
The GUI of the tracking module in Usiigaci. PCM images of a time-lapse experiment are shown in the left panel to compare with the Mask R-CNN segmented masks in the right panel. After tracking, cell tracks are listed on the right and users can verify data against PCM images and exclude bad cell tracks.

We characterized tracking performance using multiple object tracking (MOT) metrics and tracking quality measures on a triplicate 10-hr NIH/3T3 electrotaxis dataset (Table S.3). MOT metrics measure the performance of trackers based on how accurately the objects in every frame are tracked. Tracking quality can be understood more intuitively by classifying individual cell tracks in tracking quality measures. Detailed definition of tracking performance is discussed in the supplementary section S1.7.

The MOT performance of Usiigaci with or without manual verification is benchmarked against manual tracking as shown in Table 2 [40, 41]. In manual tracking, the multiple object tracking precision (MOTP) and multiple object tracking accuracy (MOTA) are arbitrarily defined as 1. A total of 4520 events are identified, summed from all frames. After tracking by the Usiigaci tracker, 4470 events are identified with MOTA of 91.9%. By imposing the *select complete track* criterion, events belonging to invalid tracks (Fig. S.6 B-H) are easily removed. The MOTPs describing the total error in positions of matched object-hypothesis pairs in Usiigaci before and after manual verification are 70.2% and 75.6%, which are similar to the Jaccard index in segmentation [40]. The masks of tracked cells correlate well with those by manual segmentation at pixel level, which suggests that cell movements and morphology changes can be tracked and analyzed quantitatively.

**Table 2:** Summary of multiple object tracking of NIH/3T3 electrotaxis after 10-hr under 300 V/m dcEF (31 frames). Metrics are compared among manual tracking and Usiigaci with and without the *select complete tracks* criterion and manual verification. MOTP: Multiple object tracking precision; MOTA: Multiple object tracking accuracy.

| MOT metrics | Manual | Usiigaci<br>(unverified) | Usiigaci<br>(select complete track) |
| --- | --- | --- | --- |
| Total events | 4520<sup>a</sup> | 4470 | 1736 |
| Miss events | 0 | 145 | 0 |
| Mismatch events | 0 | 70 | 0 |
| False positive events | 0 | 165 | 0 |
| MOTA | 1 | 0.919±0.01 | n/a |
| MOTP (mIoU) | 1 | 0.702±0.012<sup>b</sup> | 0.756±0.009<sup>b</sup> |
| **Tracking quality measure** |  |  |  |
| Total tracks | 155<sup>c</sup> | 291<sup>d</sup> | 61 |
| Valid single tracks | 104 | 56 | 56 |
| Interrupted single cell tracks | 0 | 21 | 0 |
| Mitosis cell tracks | 5 | 5 | 5 |
| Entering viewfield tracks | 19 | 19 | 0 |
| Loss of tracking tracks | 0 | 152 | 0 |
| Exiting viewfield tracks | 27 | 27 | 0 |
| Mismatch tracks | 0 | 2 | 0 |
| False positive tracks | 0 | 9 | 0 |
| Valid track ratio | 0.67<sup>e</sup> | 0.19<sup>f</sup> | 0.92<sup>g</sup> |

<sup>a</sup> Total objects identified by a human operator.
<sup>b</sup> Mean intersection over union ratio of all matched-object pairs in mean±standard error of mean.
<sup>c</sup> Total cell tracks identified by a human operator.
<sup>d</sup> Total cell tracks generated by Usiigaci’s tracker.
<sup>e</sup> Ratio of valid cell tracks to total cell tracks in the dataset identified by a human operator.
<sup>f</sup> Ratio of valid cell tracks to total cell tracks generated by Usiigaci’s tracker.
<sup>g</sup> Ratio of valid cell tracks to total cell tracks after the *select all tracks* criterion.

Tracking quality using the Usiigaci tracker can be understood more intuitively by classifying individual cell tracks. By manual tracking, 104 valid tracks are found among 155 total tracks. Using the Usiigaci tracker, 291 tracks are generated and many of which are erroneous due to different types of error (Fig S.6). The valid track ratio in Usiigaci is only 19.5% without manual verification. However, by the *select complete tracks* criterion, users can select only the tracks with the same ID in every frame. Valid cell tracks will be among those selected with the criterion. Users can also verify whether there are any erroneous tracks and exclude them if necessary. Five mitosis tracks exist in the remaining results and they are excluded manually. The valid tracks obtained from Usiigaci after manual verification correspond to 54% of valid tracks identified by a human operator. However, more viewfields can be analyzed to increase the number of valid tracks with the labor-saving workflow of Usiigaci.

### 3.3. Data processing module

Quantitative cellular dynamics require both accurate cell segmentation and cell tracking. After tracking, the data processing module of Usiigaci generates quantitative results of step-centric and cell-centric parameters in cell migration based on the tracking results. Visualization of cell migration is carried out automatically to generate visual representations that can be understood intuitively (Fig. S.9 in the SI document).

We further examine overall accuracy in the context of cell migration among the results segmented and tracked using various methods. Directedness is a metric to show directional cell migration. Directness is defined as the average cosine between the net trigonometric distance and electric current vector (Fig. S.1B). A group of cells migrating toward the cathode has a directedness of 1, and random migrating cells possess a directedness of 0 (Table S.1). The directedness of NIH/3T3 cells in dcEF is used to benchmark the accuracy of results tracked by various tracking methods including manual tracking in ImageJ, the track object module in Metamorph, Imaris Track, and tracking with Lineage Mapper (Figure 6 & Fig. S.10). PCM images, fluorescence images, or segmented masks from either Usiigaci, PHANTAST, Fogbank, or Deepcell are used in each tracking software accordingly. Only valid cell tracks that contains cells being tracked in every frame are analyzed. Capture rate is defined as the ratio between valid cell tracks by a certain method and valid cell tracks identified manually.

**Figure 6:**
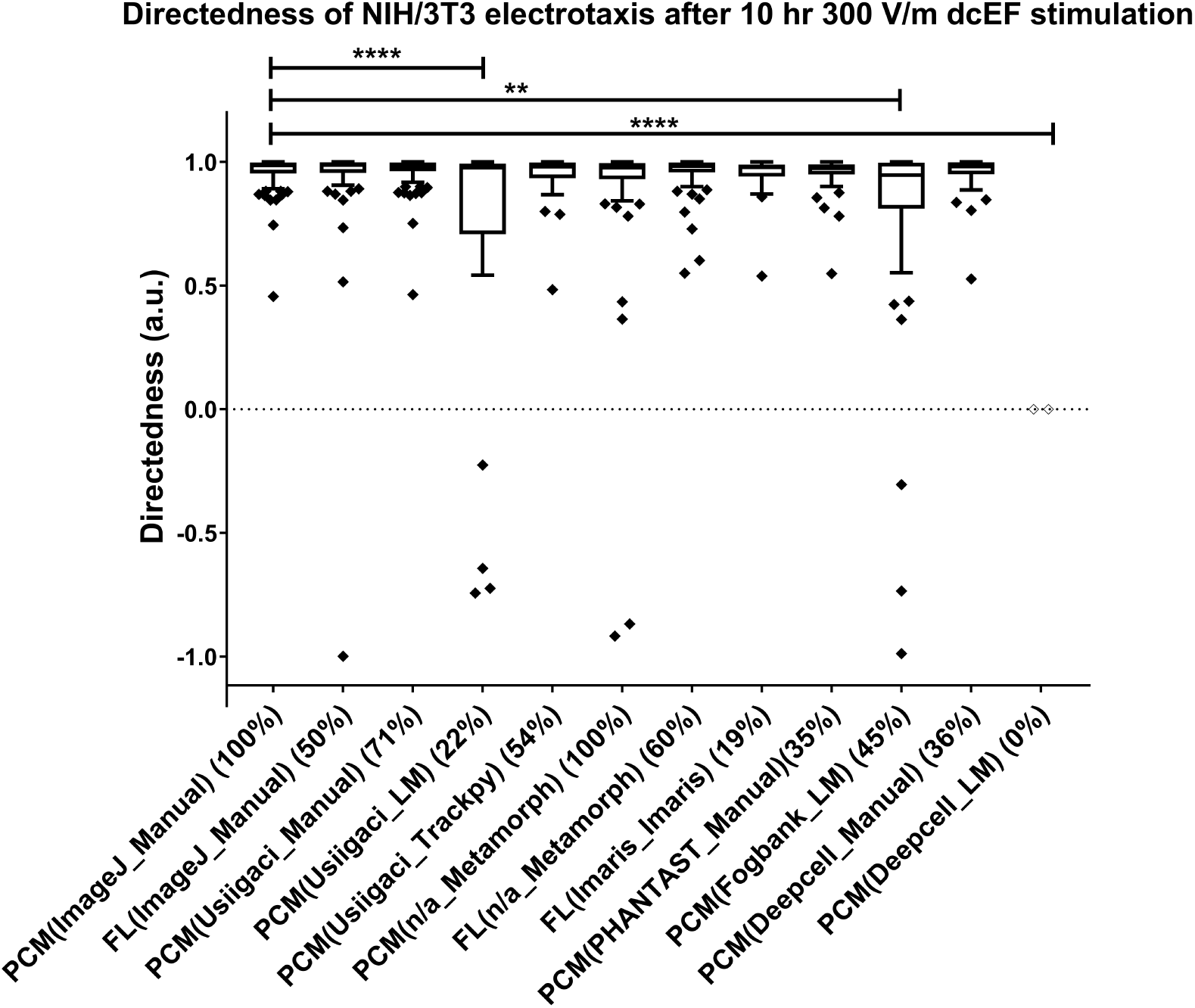
Directedness of NIH/3T3 electrotaxis after 10-hr, 300 V/m dcEF stimulation analyzed by different segmentation and tracking methods. Data and labels are arranged based on the type of images-(segmentation method tracking method) (capture rate). LM:Lineage Mapper; FL:fluorescence; PCM: phase contrast microscopy; ** denotes P<0.01; **** denotes P<0.0001.

While cell tracking in proprietary software such as Imaris and Metamorph yields results similar to the manual reference, both software packages only provide positional information about cells, while morphological information of cells is not available. Moreover, Imaris demands fluorescent labeling of cells to obtain good segmentation results (Table S.4).

Even though open-source cell tracking software, such as Lineage Mapper is available [42], segmented data may not be directly compatible with Lineage Mapper if single cells are not segmented into individual instances correctly in every frame. Because Lineage Mapper is fully automatic, a manual verification process is not available in Lineage Mapper. Imperfect segmentation results lead to erroneous tracking results and invalid tracks cannot be excluded by users. Directedness of cells segmented by Fogbank and tracked by Lineage mapper (P<0.01, Tukey’s post-hoc) differs from the manual reference. Cells segmented by Deepcell are not tracked well with Lineage Mapper (P<0.0001, Tukey’s post-hoc). Therefore, segmented results from PHANTAST and Deepcell on NIH/3T3 electrotaxis cannot yield good data by tracking with Lineage Mapper. While Usiigaci’s segmented masks can also be tracked using Lineage Mapper, only 22% of cells are tracked compared to the manual tracking reference. The results from Lineage Mapper are also significantly different compared to a manual reference (P<0.0001, one-way ANOVA), presumably due to erroneous tracking that cannot be verified manually. Misinterpretation may be made due to bad results if users do not fully grasp the inner workings of the tracking process (Figure 6).

In contrast, by segmenting and tracking with Usiigaci, 54% of cells can be automatically tracked when compared to manual tracking. Moreover, directedness and migration speed of cells analyzed by Usiigaci are comparable to the manual reference and Metamorph. Migration speed can be over-or under-estimated in Imaris or Lineage Mapper. Detailed tracking results are shown in Table S.4 in the SI document.

Usiigaci is the only automated cell tracking method that provides both cell movement and morphology change information among benchmarked software packages. With high segmentation and tracking accuracy, Usiigaci delivers quantitative cell migration analysis to biologists as an easy-to-use tool. A tutorial video of Usiigaci’s usage is provided in the supplementary information (Video S.1).

## 4. Impact and conclusions

Usiigaci offers a reliable quantitative solution for segmentation, tracking, and analysis of cell migration in two-dimensional PCM. No label or special treatment of cells is required, so that cells can be analyzed under more natural conditions. Entire outlines of cells are automatically segmented and tracked in Usiigaci, which enables biologists to analyze both movement and morphological changes in cellular dynamics in a quantitative manner that existing software cannot provide. The labor-saving workflow also alleviates the workload in comparison to the manual cell tracking method that is conventionally adopted. The manual verification function enables users to verify the tracking data and ensure data validity. The analytical capability of Usiigaci can contribute to the international effort to standardize cell migration experiments [43]. The trainable nature of the Mask R-CNN model allows Usiigaci to analyze images acquired in other bright-field microscopic techniques, and potentially for 3D cell tracking in the near future. Similar deep learning methods for biomedical image analysis are used to accomplish *in silico* labeling of cellular components instain-free images and 3D segmentation of noisy medical images [44–47]. Advances in deep learning methods for biomedical image analysis provide unique opportunities to advance biomedical discovery.

## Acknowledgements

This work is supported by JSPS KAKENHI [Grant Number JP1700362]. H.-F. Tsai and A.Q. Shen also thank Okinawa Institute of Science and Technology Graduate University (OIST) for its financial support with subsidy funding from the Cabinet Office, Government of Japan. Funders had no role in study design, data collection, the decision to publish, or preparation of the manuscript. The authors acknowledge support from the Scientific Computing and Data Analysis Section, the Community Relations Section, and the Imaging Analysis Section of OIST Graduate University. The authors also thank Matterport Inc. for their Mask R-CNN implementation source code released under the MIT license for use in part of this work. The authors thank Mr. Emanuele Martini for his open-source BW Jtrack ImageJ plugin. The authors acknowledge Ms. Tsai, Yi-Ching and Ms. Shivani Sathish from Micro/Bio/Nanofluidics Unit at OIST for assistance in preparation of illustrations in this work. The authors thank Dr. Steven Aird, OIST’s technical editor for proofreading this article.

## Supplementary Information

Supplementary information includes detailed description on cell migration experiments, microscopy protocols, annotation of training dataset, training process on the Mask R-CNN model, evaluation of multiple object tracking benchmark, and discussions on the limitation of Usiigaci. A video tutorial of Usiigaci is also attached (Video S.1).

## Required Metadata

Current code version

**Table 3:** Code metadata

| **Nr.** | **Code metadata description** | **Please fill in this column** |
| --- | --- | --- |
| C1 | Current code version | v1.0 |
| C2 | Permanent link to code/repository used for this code version | *https://github.com/oist/Usiigaci* |
| C3 | Legal Code License | MIT License |
| C4 | Code versioning system used | git |
| C5 | Software code languages, tools, and services used | Python, TensorFlow, Keras, Trackpy, NumPy, SciPy, Pandas, PyQtGraph |
| C6 | Compilation requirements, operating environments & dependencies | Ubuntu 16.04 Linux, Python3.4+, CUDA9.1, TensorFlow 1.4, Keras 2.1 |
| C7 | If available Link to developer documentation/manual | None |
| C8 | Support email for questions |  |

