## Supplementary information for "Usiigaci: Instance-aware cell tracking in stain-free phase contrast microscopy enabled by machine learning"

### S1 Materials and methods

#### S1.1 Cell culture and maintenance

Mouse embryonic NIH/3T3 fibroblasts (CRL-1658, American Type Culture Collection, ATCC, USA) and T98G glioblastoma cells (CRL-1690, ATCC, USA) were cultured and maintained according to guidelines provided by ATCC. NIH/3T3 cells were cultured in Dulbecco's modified Eagle's medium (12800017, DMEM, Gibco, USA) supplemented with 2.2 g/L $NaHCO_3$ and 10% iron-fortified calf serum (CS, Sigma-Aldrich, USA). T98G cells were cultured in minimum essential medium alpha (12000022, MEM$\alpha$, Gibco, USA) supplemented with 2.2 g/L $NaHCO_3$ and 10% fetal bovine serum (FBS, Gibco, USA). Both cell lines were cultured in a humidified 5% $CO_2$ atmosphere (MCO-18AIC, Sanyo, Japan) and subcultured whenever confluency reached 80%. Cells from passages 5 to 15 were used in the migration study. Fluorescently labeled cells were prepared by incubating $5 \times 10^6$ cells with 1 $\mu$M of CellTracker Green CMFDA dye (ThermoFisher, USA) in 1X Dulbecco's phosphate buffered saline (D-PBS, Wako Inc., Japan) for 30 minutes at 37°C according to the protocol provided by the manufacturer. Cells were washed once in D-PBS and seeded into microfluidic chips for microscopy.

#### S1.2 Electrotaxis experiment

A microfluidic electrotaxis chip containing two channels with L×W×H of $30 \times 3 \times 0.07$ mm was composed of a top Poly(methyl methacrylate) (PMMA) chip, a piece of double-sided tape (PET8018PT, 3M, USA) cut with a $CO_2$ laser scriber (VLS3.50, Universal Laser Systems, USA) to define the microfluidic channels, and a coverglass substrate (0107242, No. 1.5 high precision, Marienfeld, Germany) (Fig. S.1.A). The PMMA microfluidic chip was fabricated by patterning inlets, outlets, and fluidic connections on PMMA sheets using a $CO_2$ laser and bonded thermally as described previously [1]. Adapters with M6 threads (SPC-M6-C, Nabeya, Japan) were glued onto the PMMA microfluidic chip using a UV adhesive (3301, Loctite, USA). The coverglass substrate was joined with the PMMA microfluidic chip using the cut double-sided tape after UV disinfection of all surfaces.

Prior to seeding the cells, the coverglass was coated with 100 $\mu$g/ml poly-D-lysine (P6407, Sigma-Aldrich, USA) or Geltrex (A1569601, Gibco, USA) for NIH/3T3 cells and T98G glioma cells for 2 hours at 37°C. $7.5 \times 10^5$ cells/ml were seeded into each channel and allowed to adhere at 37°C. Afterward, the tubing for infusing media and salt bridges were connected to the microfluidic chip containing adhered cells by fittings (IDEX, USA). The chips were moved onto a Nikon Ti-E microscope equipped with an in-house transparent heater made of indium-tin-oxide (ITO) glass. Media were steadily infused at 20 $\mu$L/hr using a two-channel syringe pump (YSP-202, YMC, Japan) and a stable 37±0.1°C temperature was provided using a heater to culture the cells. A stable electrical current (101.5 $\mu$A for 300 V/m EF) was applied with a sourcemeter unit (SMU, 2410, Keithley, USA) through salt bridges (1.5% agarose in 1X D-PBS) with a pair of silver/silver chloride electrodes in 1X PBS. Silver/silver chloride electrodes were fabricated by electroplating a pure silver sheet (0.1 mm-thick, Nilaco, Japan) in 1 N HCl (Sigma-Aldrich, USA) after being briefly etched in 20% nitric acid (Sigma-Aldrich, USA) [2]. To quantitatively analyze cell migration in electrotaxis experiments (Fig. S.1.B), cell-centric and step-centric parameters were extracted and analyzed (Table S.1).

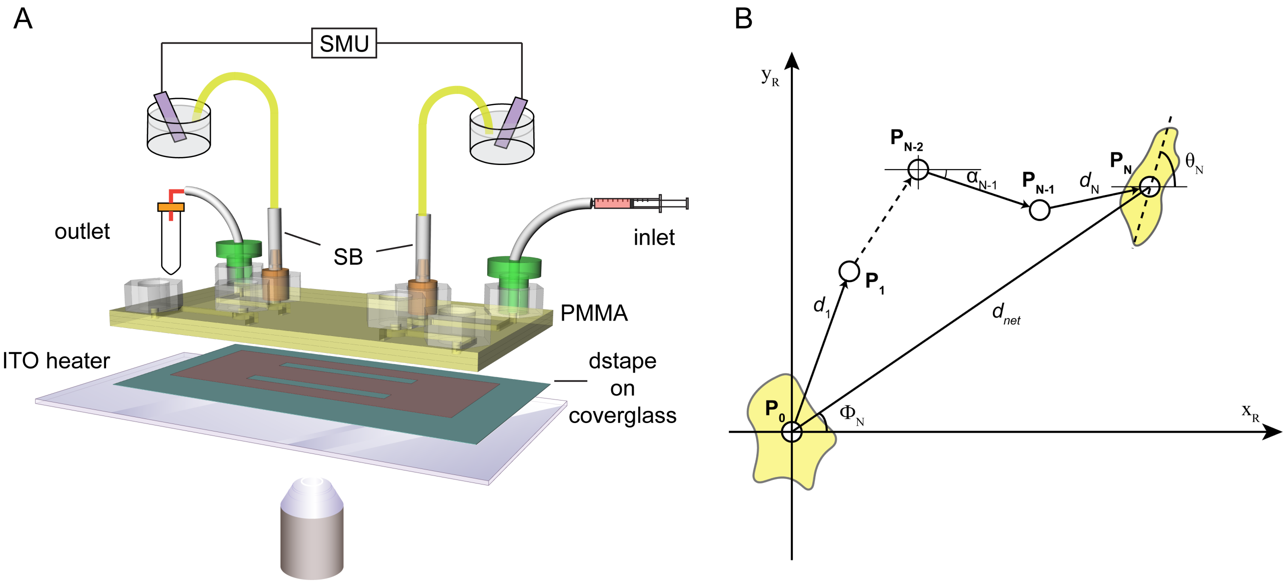


Fig. S.1: (A) The setup diagram of an electrotaxis experiment using a microfluidic chip. SB: salt bridges; SMU: sourcemeter unit. (B) The single-cell migration parameters extracted from the electrotaxis experiments.

Table. S.1: Step-centric and cell-centric parameters for describing single-cell migration.

| | |
|---|---|
| **Step-centric features** | |
| Instantaneous displacement | $d_i = \sqrt{(x_i - x_{i-1})^2 + (y_i - y_{i-1})^2}$ |
| Instantaneous speed | $s_i = d(p_{i-1}, p_i)/\Delta t$ |
| Turning angle | $\alpha_i = \tan^{-1}\frac{(y_i - y_{i-1})}{(x_i - x_{i-1})}$ |
| Directional autocorrelation | $dir - aut_i = \cos(\alpha_i - \alpha_{i-1})$ |
| **Cell-centric features** | |
| Cumulative distance | $d_{total} = \sum_{i=1}^{N} d(p_{i-1}, p_i)$ |
| Euclidean (net trigonometric) distance | $d_{net} = d(p_0, p_N) = \sqrt{(x_N - x_0)^2 + (y_N - y_0)^2}$ |
| Euclidean velocity | $\bar{v}_{net} = \frac{d_{net}}{t_{elapsed}}$ |
| End-point directionality ratio | $ep\_dr = \frac{d_{net}}{d_{tot}}$ |
| Orientation | $\text{Index}_{\text{orientation}} = \sum_{i=1}^{N} \frac{\cos 2\theta_i}{N}$ |
| Directedness | $\text{Index}_{\text{directedness}} = \sum_{i=1}^{N} \frac{\cos \Phi_i}{N} = \sum_{i=1}^{N} \frac{(x_i - x_1)}{d_{net} \times N} = \frac{1}{N} \sum_{i=1}^{N} \frac{(x_i - x_1)}{\sqrt{(x_i - x_1)^2 + (y_i - y_1)^2}}$ |

#### S1.3 Cell microscopy

A Nikon Ti-E microscope with Perfect Focus System and motorized XY stage was used to perform all microscopy experiments. A 10X phase contrast objective and an intermediate magnification of 1.5X were used for phase contrast and epifluorescence imaging. Images were captured with a sCMOS camera with 2×2 binning (Orca Flash 4.0, Hamamatsu, Japan) at 20-minute interval in NIS Element software (Nikon, Japan). The spatial resolution at this setting was 0.87 $\mu$m/pixel. For fluorescence imaging of CellTracker Green, a FITC-channel filter set was used (FITC-3540C-NTE-ZERO, Semrock, USA) with an Intensi-light fiber mercury lamp (Nikon, Japan) as the excitation light source. The Perfect Focus System was used for all time-lapse experiments and all experiments were performed in triplicate. After each experiment, images were exported from NIS element as tiff files and automatically organized according to the XY positions using an in-house Python script included in the Usiigaci's source code.

#### S1.4 Training dataset annotation

Fifty phase contrast microimages of T98G and NIH/3T3 cells were manually segmented in Fiji ImageJ [3] with the aid of a drawing tablet (Cintiq pro 16, Wacom, Japan) (Fig. S.2). Briefly, the boundary of each cell was traced manually with the freehand tool in ImageJ and saved into the ROI manager. Indexed masks were created from the manually traced ROIs by using the ROImap function in the ImageJ plugin LOCI [4]. A unique index for each ROI, *i.e.,* for each cell, is essential for instance-aware segmentation and downstream tracking. Phase contrast images were used in tiff format without modification. Forty-five sets of images were used as the training dataset and the rest as the validation dataset.

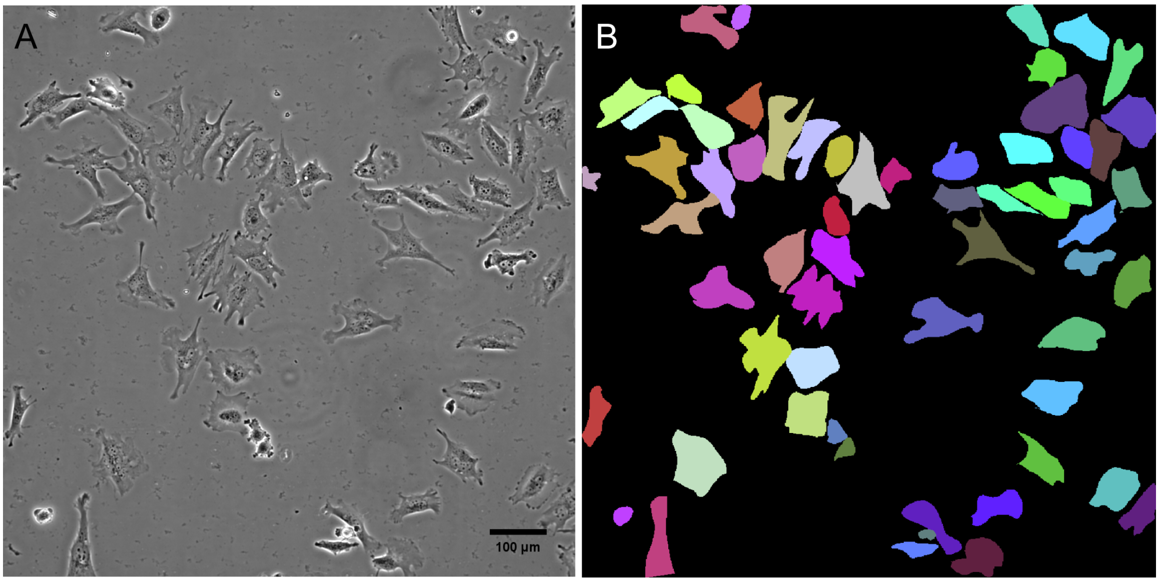


Fig. S.2: An example of the training data. (A) Phase contrast image of T98G cells on Geltrex-coated substrate. (B) Cell outlines were manually segmented and saved into an 8-bit indexed mask. Each color represents a unique set of cells.

#### S1.5 Training and deployment of Usiigaci for generation of instance-aware masks

The Mask R-CNN model (Fig. S.3) of Usiigaci was trained using our annotated training dataset on top of a pretrained Mask R-CNN model from Matterport Inc. The pretrained Mask R-CNN model was trained with the Microsoft COCO dataset [5]. Additional training of 100 epochs on the feature pyramid network headers and 500 epochs on all ResNet-101 layers were carried out under a Ubuntu 16.04 environment with Python 3.4, TensorFlow 1.4, Keras 2.1.2, and CUDA 9.1 on an Alienware 15 laptop equipped with a GTX1070 8GB graphics processing unit (GPU) or a NVIDIA GTX1080Ti 11GB GPU (Manli, Hong Kong) mounted on an alienware amplifier (Dell, USA).

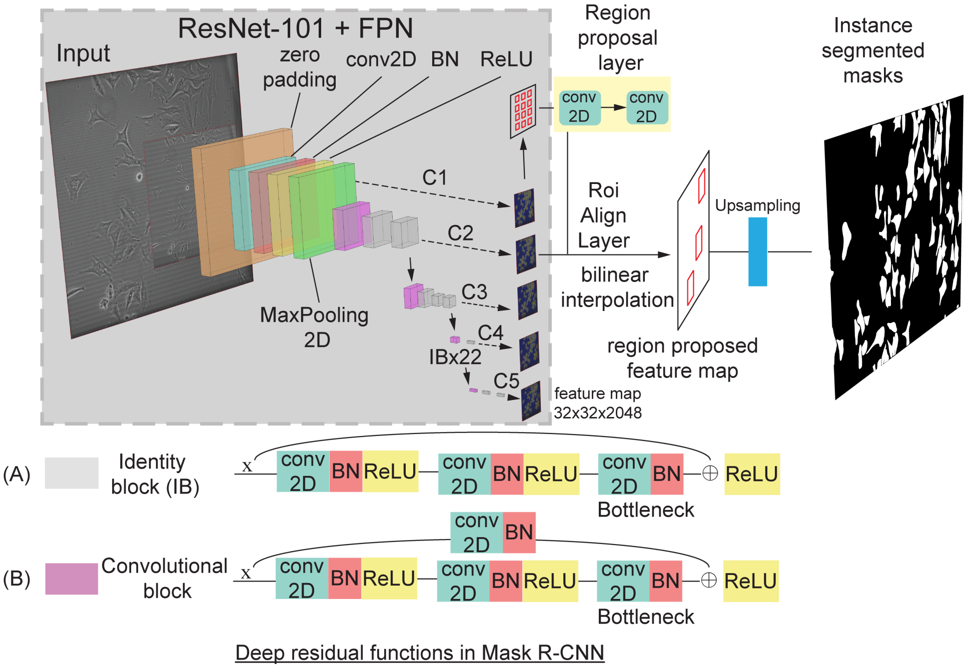


Fig. S.3: Architecture of Mask R-CNN module in Usiigaci. The backbone of Mask R-CNN model is a combination of 101-layer deep residual network (Resnet-101) [6] and feature pyramid network (FPN) [7]. The ResNet-101 is a deep residual network composed by two types of deep residual functions: (A) identity block and (B) convolutional block.

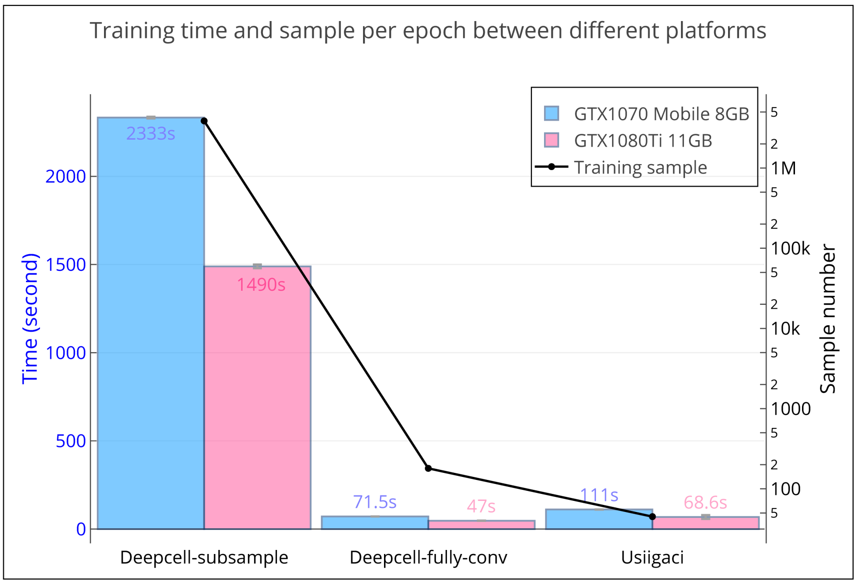


Fig. S.4: Comparison of training time and sample number per epoch using Usiigaci and Deepcell in subsample mode and in fully convolutional mode.

When morphologies of the cells or microscopy optical configurations change significantly, retraining the neural network may be necessary for optimal results. A training script was provided as part of the source code to allow users to train new Mask R-CNN models based on their own data for best performance after proper training data annotation as described previously. The best model was selected by observing accuracy and loss during the validation step in training. In Usiigaci's Mask R-CNN segmentation module, it is essentially a two-class segmentation system, background and whole-cell outlines. Several hyper-parameters for training can be adjusted by users, such as neural network backbone configuration, threshold for region proposals network, and number of anchors for regional proposal networks.

In addition, to compare with Deepcell, a state-of-the-art machine learning-based cell segmentation software, a TensorFlow implementation of Deepcell from the original authors was modified to use only the phase contrast channel for segmentation [8]. The modified Deepcell was trained from scratch using the aforementioned 50 sets of manually annotated training data. The sample size and training time per epoch were compared between Usiigaci's Mask R-CNN module and Deepcell in subsample mode and fully convolutional mode (Fig. S.4). In the subsample mode of Deepcell, training data were split into small batches based on the set window size to increase arbitrary training samples, and the computation time required to complete an epoch increased. A moderate computational performance improvement was observed when using a GPU that had more CUDA cores (GTX1080Ti). Usiigaci's Mask R-CNN model and Deepcell in fully convolutional mode used full-size images for training with simple image augmentation performed at the start of end epoch. Therefore, the time required to complete an epoch was shorter compared to Deepcell in subsample mode, but more epochs were needed to increase the accuracy of the results. Training of Mask R-CNN took only slightly more time than Deepcell despite more layers and trainable parameters in the Mask R-CNN model because the Mask R-CNN had already been trained for edge detection with the Microsoft COCO dataset (101 layers, 64.7 million parameters compared to 25 layers and 0.46 million parameters in Deepcell).

In general, increasing the number of training data will increase the accuracy of the CNN, but our work demonstrated that a pretrained Mask R-CNN model could be trained using a small training dataset to achieve reliable segmentation with mean intersection over union (mIoU) of 0.72 in an example dataset that the R-CNN model was not exposed to during the training (Figure 4 in the main manuscript). The performance of Mask R-CNN surpassed several segmentation software we benchmarked. The fact that a small training dataset can fulfill the need for reliable segmentation, this can be appealing to end users who do not possess extensive computational or experimental resources to generate large training datasets.

Deployment was performed under the same environment as training with GPU acceleration of either a NVIDIA GTX1070 8GB GPU or a NVIDIA GTX1080Ti 11GB GPU. The inference python script for deployment allowed users to specify a folder path with nested folders, and Usiigaci segmented all images in each nested folder and created 8-bit indexed mask images with each cell possessing a unique identifier. Segmentation with multiple trained models and averaging among the results can be specified in the Python script if necessary.

#### S1.6 Cell segmentation accuracy evaluation

Phase contrast microscopy images of NIH/3T3 mouse fibroblast electrotaxis under 300 V/m direct current electric field (dcEF) for 10 hr in the microfluidic chip were collected (Fig. S.1.A). The experiments were performed in triplicates. Segmentation similarity of Usiigaci was compared with existing state-of-the-art free software for cell segmentation, including fluorescent thresholding, PHANTAST [9], Fogbank [10], and Deepcell [8]. The ImageJ versions of Fogbank and PHANTAST were used for segmentation similarity evaluation. Segmentation similarity was compared against human segmentation (manual reference) by segmentation evaluation metrics (Fig. S.5 & Table S.2). True positive (TP), true negative (TN), false positive (FP), and false negative (FN) metrics were extracted by comparing the manual reference and segmentation results. Jaccard index and F1 scores were calculated for each segmentation method.

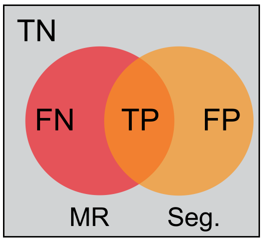


Fig. S.5: Segmentation performance evaluation. MR: Manual reference; Seg.: Segmentation results; FN: False negative; TP: True positivie; FP: False positive; TN: True negative.

Table. S.2: Segmentation evaluation metrics to describe the similiarity between manual reference (MR) and segmentation (Seg.) results.

| **Segmentation evaluation metrics** | |
|---|---|
| True positive | TP $= MR \cap Seg.$ |
| True negative | TN $= MR \cup Seg.$ |
| False positive | FP $= Seg. - MR$ |
| False negative | FN $= MR - Seg.$ |
| Jaccard index (IoU) | $\text{Index}_{\text{Jaccard}} = \frac{TP}{FN+TP+FP}$ |
| F1 score | $\text{F1} = \frac{2TP}{2TP+FP+FN}$ |
| Precision | $\text{Precision} = \frac{TP}{TP+FP}$ |
| Recall | $\text{Recall} = \frac{TP}{TP+FN}$ |
| Specificity | $\text{Specificity} = \frac{TN}{TN+FP}$ |
| Accuracy | $\text{Accuracy} = \frac{TP+TN}{TP+TN+FP+FN}$ |

#### S1.7 Cell tracking accuracy evaluation

In each experimental dataset, different tracking results could be identified and categorized to describe the performance of a tracker using multiple object tracking metrics (Fig. S.6 & Table S.3) [11, 12]. In this dataset, valid single cell migration tracks were the data of interest. However, due to segmentation errors and tracking errors, the data also contained several invalid tracking results, such as interrupted tracks, mitosis tracks, loss of tracking, objects entering or exiting viewfield, mismatch tracks, and false positive tracks (Fig. S.6). Interrupted tracks were tracks that were tracked correctly most of the time (>80% of time) but missing partial information. Mitosis tracks contained cells that had undergone mitosis, thus could longer represent true single cell migration. Tracks containing cells that were lost, entering, or exiting viewfield early in the experiment could not be analyzed either. Mismatch and false positive cell tracks must also be excluded.

Using the multiple object tracking metrics in computer vision, we used the multiple object tracking precision (MOTP) and multiple object tracking accuracy (MOTA) to benchmark Usiigaci's tracker performance against manual tracking by a human operator (first author). The multiple object tracking precision (MOTP) described the total error in estimated positions for matched object-hypothesis pairs over all frames, averaged by the total number of matches recognized [11]. MOTP showed the ability of the tracker to estimate precise object positions. We described the MOTP by means of average pixel level precision by comparing the polygon shape of each object between the manual reference and the segmentation result (mean intersection over union). The multiple object tracking accuracy (MOTA) accounted for all object configuration errors made by the tracker, including missed events, mismatch events, and false positive events. The detailed definition is shown in Table S.3.

In addition, we used intuitive tracking quality measures to describe the amount of valid data based on validity of cell tracks considering the uniqueness of cell migration data (Table S.3). Usiigaci tracker featured a graphical user interface (GUI) to allow users to manually verify tracking data. After tracking, valid cell tracks

Table. S.3: Multiple object tracking performance metrics to evaluate the precision and accuracy of the tracking results.

| **Multiple object tracking metrics by events [11, 12]** | |
|---|---|
| Miss events | $\overline{m} = \frac{\sum_t m_t}{\sum_t g_t}$ |
| Mismatch events | $\overline{mme} = \frac{\sum_t mme_t}{\sum_t g_t}$ |
| False positive events | $\overline{fp} = \frac{\sum_t fp_t}{\sum_t g_t}$ |
| Multiple object tracking accuracy (MOTA) | $\text{MOTA} = 1 - \frac{\sum_t (m_t + mme_t + fp_t)}{\sum_t g_t}$ |
| Multiple object tracking precision (MOTP) | $\text{MOTP} = \frac{\sum_{i,t} \frac{TP_t^i}{FN_t^i + TP_t^i + FP_t^i}}{\sum_t c_t}$ |
| **Single cell migration tracking quality measure (this work)** | |
| Valid single cell track (Fig. S.6.A) | A single cell is tracked correctly in every frame. |
| Interrupted single cell track (Fig. S.6.B) | A single cell is tracked correctly most of the time (>80%). |
| Mitosis cell track (Fig. S.6.C) | A single cell underwent mitosis. |
| Entering viewfield track (Fig. S.6.D) | A single cell enters viewfield during the experiment. |
| Loss of tracking track (Fig. S.6.E) | A single cell tracking is lost during the experiment. |
| Exiting viewfield track (Fig. S.6.F) | A single cell exits viewfield during the experiment. |
| Mismatch track (Fig. S.6.G) | Two cell tracks with ID switched erroneously during the experiment. |
| False positive track (Fig. S.6.H) | A single cell falsely tracked. |
| Valid track ratio | Ratio of valid single cell tracks to all tracks identified by tracker |

were mixed with invalid tracks, but valid cell tracks could be selected by a simple criterion, *select complete tracks*, in which only cell tracks with objects that were tracked in every frames in the experiment were selected. The valid cell tracks were then inspected by users through cross-referencing with raw image sequences to ensure the validity of the tracking result (Figure 5 in the main manuscript).

Furthermore, we benchmarked the overall performance of segmentation and tracking under the single cell migration analysis context (Fig. S.6 & Table S.4). Various segmentation and tracking methods including Usiigaci's segmentation and tracking module, Lineage Mapper [13], ImarisTrack module of Imaris (v8.4, Bitplane Inc, UK), and the track object module of Metamorph (Molecular Devices, USA) were benchmarked against manual tracking in ImageJ. Segmentation results from Usiigaci, PHANTAST, Fogbank, and Deepcell as well as phase contrast raw images, or fluorescence images were input as data into each software package as needed. An ImageJ plugin (BW_Jtrack) developed by Mr. Emanuele Martini can assist ImageJ tracking and increase manual tracking speed. Cell-centric parameters of cell migration were used to validate tracking accuracy compared to manual tracking. All data were represented as the mean ± 95% confidence interval unless otherwise noted. Statistical comparison of results from different tracking methods was done using one-way analysis of variance (ANOVA) with Tukey's post-hoc multiple-comparison test. The significance level to reject a null hypothesis between two datasets was set at 0.05. A p-value (P, the probability for a true null hypothesis) less than 0.05 represented statistical significance with corresponding 95% confidence level.

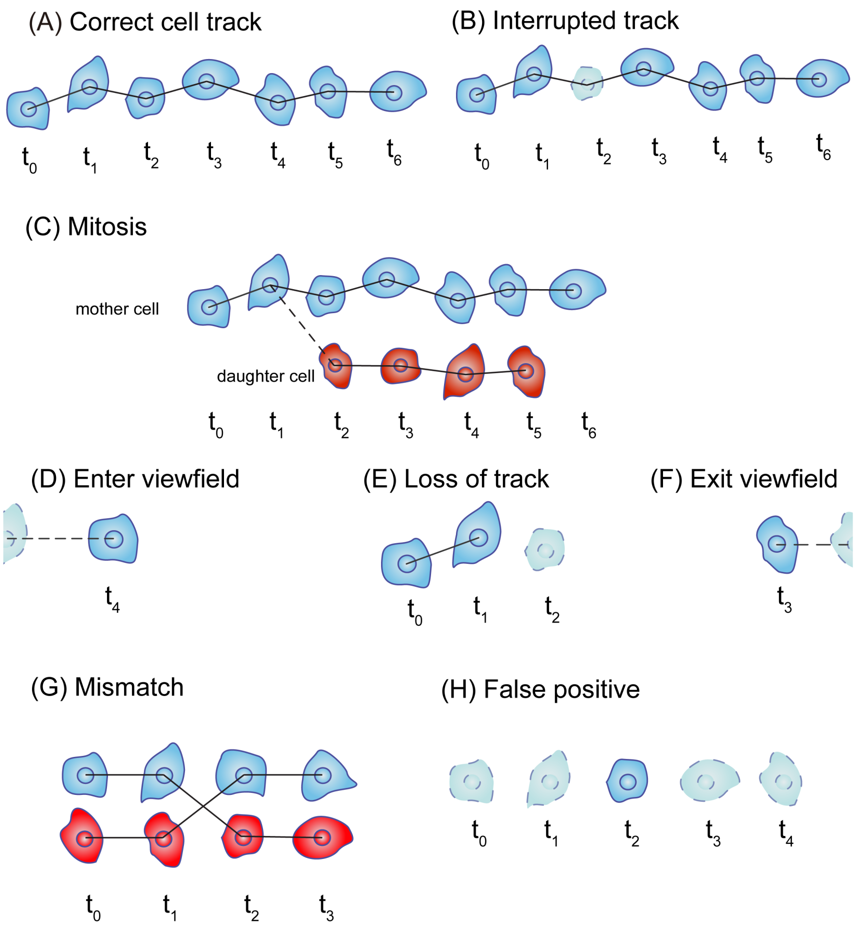


Fig. S.6: Types of cell tracking results. (A) Valid cell tracks that are identified consistently and tracked correctly in every frame throughout one time lapse experiment. (B) Interrupted cell tracks that are tracked correctly (>80% of the time) from start to end in one time lapse experiment but the data is missing in one or a few frames within the experiment. (C) Mitosis cell tracks in which cells undergo mitosis and split into daughter cells. The tracking can no longer represent true single cell migration. (D) Cell tracks entering the viewfield in the middle of the time lapse experiment. (E) Cell tracking fails in the middle of the time lapse experiment. (F) Cell tracks exiting the viewfield in the middle of the time lapse experiment. (G) Cell tracks that are linked incorrectly in tracking. (H) False positive cell tracks identified by tracker due to primarily segmentation artifacts. Tracking results in (B–F) are considered as invalid tracks.

### S2 Results and Discussion

#### S2.1 Mask R-CNN realizes fast and highly accurate whole-cell instance-aware segmentation

Segmentation results from Usiigaci are highly accurate. Fig. S.7 shows a segmentation comparison between the manual reference, fluorescence-thresholded results, Deepcell results, and Usiigaci results. In cell migration, cells in mesenchymal migration often have thin protruding cellular structures powered by the cytoskeleton to effect mechanical displacement, such as blebs or lamellipodia [14]. These thin structures have very low contrast even under PCM. Segmentation results from Usiigaci closely resemble those of manual segmentation, while both fluorescence-thresholding and Deepcell have weak segmentation on weakly contrasted cell features such as lamellipodia. Correct segmentation of the entire cell outline enables researchers to not only track cell movement, but also monitor cell morphology.

One of the drawbacks of machine learning is the necessity of powerful computing resources for training as well as deployment of such tools. By taking advantage of the Mask R-CNN model, segmentation speed is comparable to that of conventional computer vision methods such as PHANTAST and Fogbank. The segmentation speed of Usiigaci is three times faster than that of Deepcell. One 1024×1022 image can be segmented into instance-aware masks in 1.5 seconds on a standard laptop equipped with a CUDA-compatible GPU (Fig. S.8). Averaging

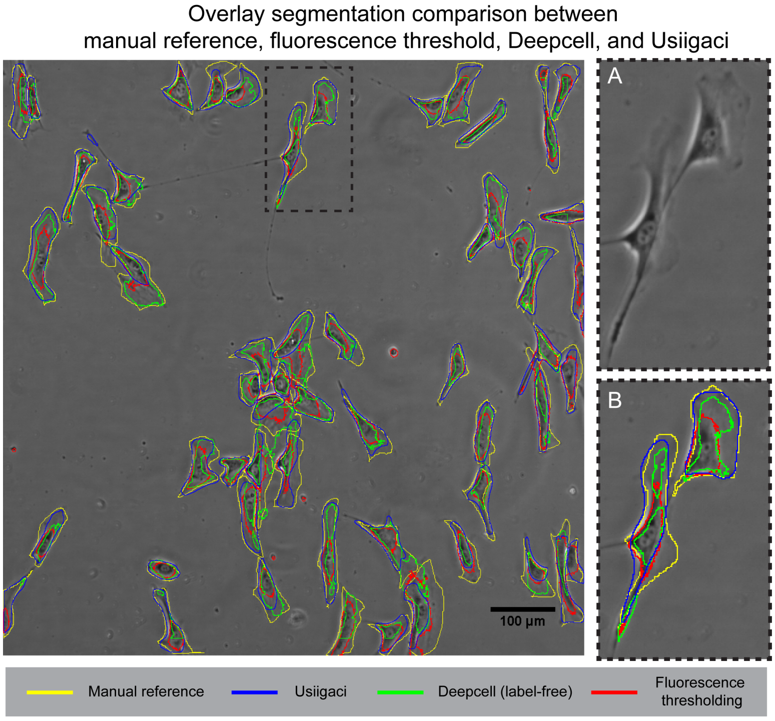


Fig. S.7: Segmentation comparison between manual segmentation as a reference (yellow), fluorescence threshold (red), Deepcell (green) and Usiigaci (blue). The magnified region of the PCM image is shown in inset A. Cells in mesenchymal migration undergo mechanical displacement using lamellipodia [14], the contrast of which under PCM is very low, making it difficult to segment the structure without labelling cell structures. Outlines of segmentation results from the three methods are overlaid in inset B. Overall segmentation accuracy on whole-cell outline by Usiigaci is more superior and resembles more closely to that of the manual segmentation reference.

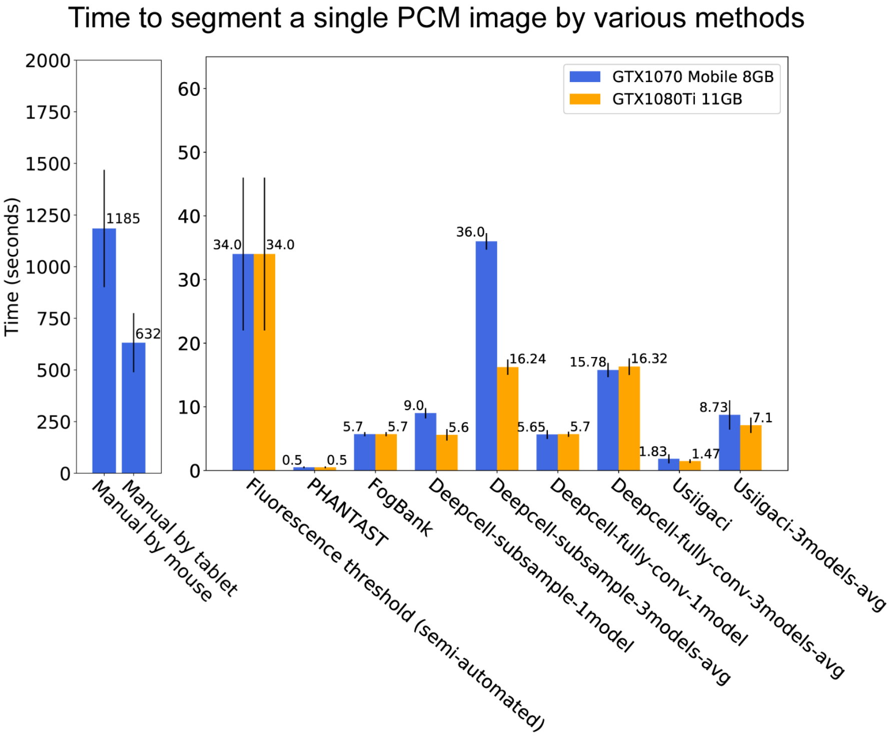


Fig. S.8: The segmentation time of one phase contrast microscopy (PCM) image acquired by various methods. Data represent the mean $\pm$ the standard error of the mean among three experiments. Note that the time scale to manually segment cell outlines is different from other methods. Blue bars indicate data processed using an on-board GPU in the Alienware laptop and orange bars indicate data processed using a desktop GPU.

segmentation results from three different model weights can still be completed in 7 seconds per image. Usiigaci's segmentation module based on Mask R-CNN model provides speed improvement of orders of magnitude over manual outline segmentation.

#### S2.2 Automated tracking with manual verification ensures reliable cell tracking results

The data processing module of Usiigaci could import data output by ImageJ, Usiigaci tracker, Imaris, or Metamorph. Step-centric and cell-centric parameters to quantify cell migration were computed automatically (Table S.1). Results were exported into spreadsheets and could be reanalyzed in other statistical software. Vsualizations of cell migration were also automatically generated by the data processing module (Fig. S.9). The module was written in Python syntax that could be easily reused if particular functionalities were needed. Usiigaci provides researchers a tool to analyze cell migration quantitatively and can potentially collect important information to validate computational cell migration models [15].

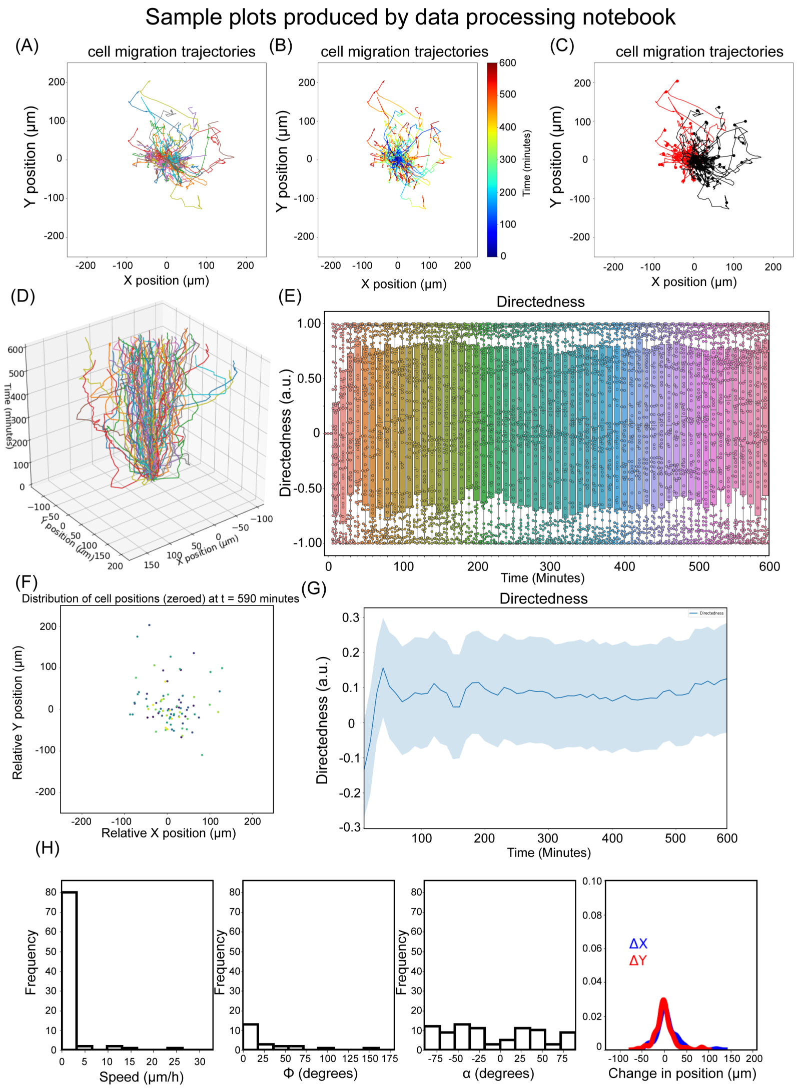


Fig. S.9: Visualization examples of NIH/3T3 random cell migration generated automatically by the data analysis module. (A) Cell migration trajectory plot with each track represented by a color. (B) Cell migration trajectory plot with cell tracks colored by time. (C) Cell migration trajectory plot with coloring depending on the displacement of cells in x direction. (D) 3D representation of cell migration trajectory. (E) Box plots of cell migration directedness versus time. (F) Cell migration represented in a scatter plot. (G) Time series plot of directedness in a time lapse experiment. (H) Frequency histogram plots of cell migration speed, orientation, turn angle, and change in positions.

Quantitative cellular dynamics analysis requires both accurate cell segmentation and cell tracking. We benchmarked the accuracy of cell segmentation and tracking under the context of single cell migration analysis with the dataset of NIH/3T3 electrotaxis under 300 V/m dcEF in a 10-hour time-lapse experiment. NIH/3T3 has demonstrated classical cathodal migration and perpendicular alignment, which is visible by the increase in directedness and the decrease in orientation (Fig. S.10). The performance of Usiigaci's Mask R-CNN segmentation and Trackpy-based tracker benchmarked against manual tracking in ImageJ, Imaris, Metamorph, Lineage mapper are shown in Table S.4.

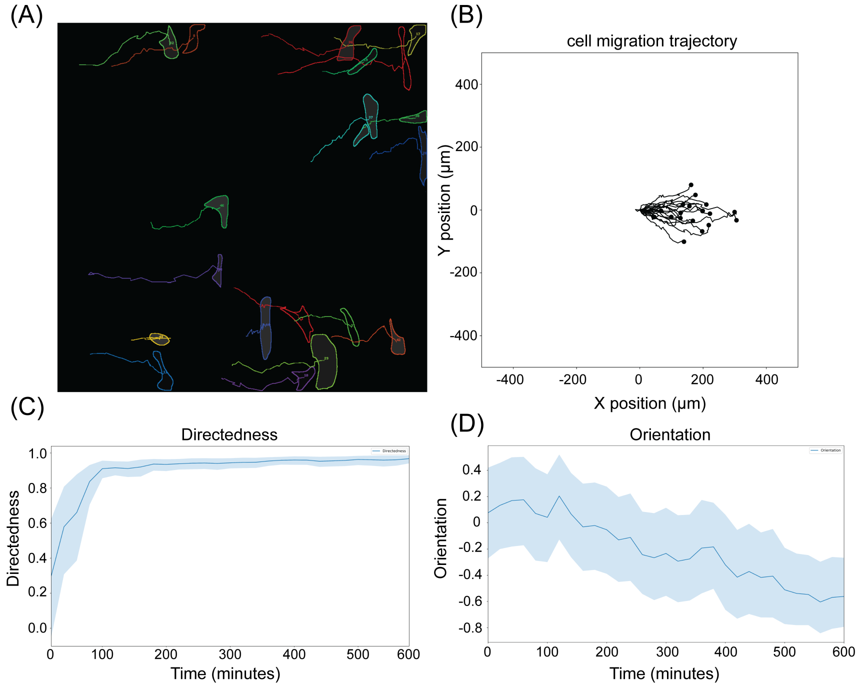


Fig. S.10: Visualization of valid tracks of NIH/3T3 10-hr electrotaxis in 300 V/m dcEF. The cathode is on the right side of the viewfield. (A) Tracked results from Usiigaci tracker; (B) Cell migration trajectory plot; (C) Directedness of cell migration; (D) Orientation of cell migration.

Usiigaci is the only all-in-one solution for automated cell tracking with manual verification functionality that provides cell movement and morphology change information. Quantitative analysis of cellular dynamics performed using Usiigaci offers unique advantages for quantitative biology studies.

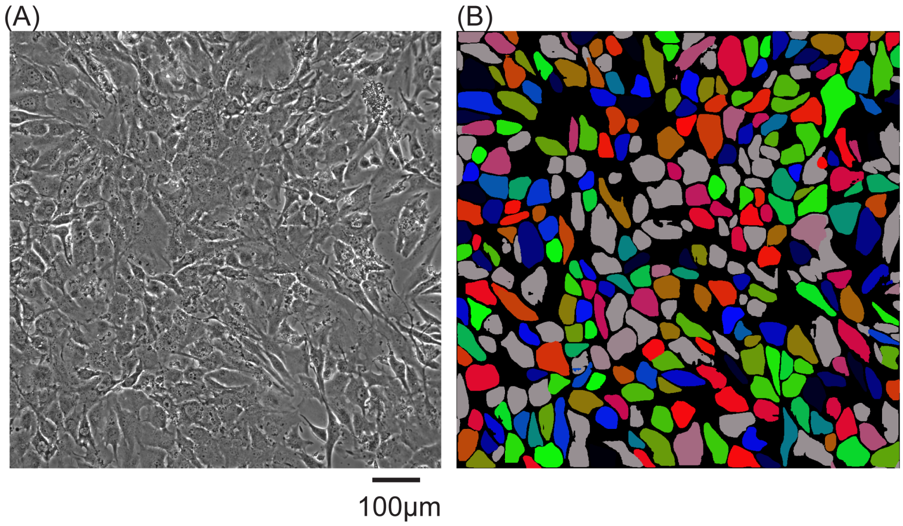


Fig. S.11: Segmentation results of confluent T98G glioma cells. (A) PCM image of confluent T98G glioma cells. (B) The corresponding segmentation result.

Table. S.4: Tracking results of NIH/3T3 electrotaxis using different segmentation and different tracking methods. Abbreviations: Ch: channel; PCM: phase contrast microscopy; FL: fluorescence; sem:standard error of mean.

| NIH/3T3 10hr 300mV/mm in triplicate | | Ch | Single cell tracks | Capture percentage | Directedness | sem | Speed ($\mu$m/hr) | sem | Orientation 0hr | sem | Orientation 10hr | sem |
|---|---|---|---|---|---|---|---|---|---|---|---|---|
| Outline tracking | manual | PCM | 104 | 100% | 0.95 | 0.02 | 18.65 | 0.69 | -0.08 | 0.08 | -0.76 | 0.03 |
| Outline tracking | manual | FL | 52 | 50% | 0.92 | 0.04 | 18.51 | 1.19 | -0.12 | 0.11 | -0.75 | 0.06 |
| Imaris | automated | FL | 20 | 19% | 0.94 | 0.02 | 21.73 | 2.09 | n/a | n/a | n/a | n/a |
| Metamorph | automated | PCM | 104 | 100% | 0.92 | 0.04 | 18.32 | 0.73 | n/a | n/a | n/a | n/a |
| Metamorph | automated | FL | 62 | 60% | 0.95 | 0.01 | 18.15 | 1.03 | n/a | n/a | n/a | n/a |
| Fogbank (Lineage Mapper) | automated | PCM | 47 | 45% | 0.79 | 0.06 | 11.37 | 3.32 | n/a | n/a | n/a | n/a |
| PHANTAST (ImageJ) | manual | PCM | 36 | 35% | 0.95 | 0.01 | 16.65 | 1.35 | -0.08 | 0.11 | -0.59 | 0.10 |
| Deepcell (ImageJ) | manual | PCM | 37 | 36% | 0.95 | 0.01 | 18.68 | 1.19 | -0.14 | 0.12 | -0.82 | 0.05 |
| Deepcell (Lineage Mapper) | automated | PCM | 0 | 0% | n/a | n/a | n/a | n/a | n/a | n/a | n/a | n/a |
| Usiigaci (ImageJ) | semi automated | PCM | 74 | 71% | 0.96 | 0.00 | 18.34 | 0.82 | -0.04 | 0.09 | -0.74 | 0.05 |
| Usiigaci (Lineage Mapper) | automated | PCM | 23 | 22% | 0.67 | 0.13 | 4.28 | 0.64 | n/a | n/a | n/a | n/a |
| Usiigaci (Trackpy) | automated | PCM | 56 | 54% | 0.92 | 0.05 | 18.19 | 1.38 | -0.09 | 0.10 | -0.80 | 0.04 |

##### S2.3 Limitation of Usiigaci

We have shown that Usiigaci's instance-level segmentation can provide reliable cell outline segmentation and tracking with improved speed. However, if the cell morphology differs significantly than that in the training data, the neural network may not perform as well as expected.

A microscopy photo of confluent T98G glioma cells and its corresponding Usiigaci segmentation are shown in Fig. S.11. The cell boundaries of confluent cell layer may be obscured and become less sharp in comparison to those seen in single cell microscopy images. Lack of clear cell boundaries makes it difficult to be recognized by the neural network. Therefore, the Mask R-CNN neural network is limited by the training data. Preparation of proper training data may be necessary for end users to obtain optimal results if cell morphology and microscopy optical configuration are significantly different than our current setup.
